# House sparrows do not exhibit a preference for the scent of potential partners with different MHC-I allele numbers and genetic distances

**DOI:** 10.1101/2022.01.11.475834

**Authors:** Luisa Amo, Guillermo Amo de Paz, Johanna Kabbert, Annie Machordom

## Abstract

MHC genes play a fundamental role in immune recognition of pathogens and parasites. Therefore, females may increase offspring heterozygosity and genetic diversity by selecting MHC genetically compatible or heterozygous males. In birds, several studies suggest that MHC genes play a role in mate choice, and recent evidence suggest that olfaction may play a role in such discrimination. Previous studies indicated that house sparrow females with low allelic diversity prefer males with higher diversity in MHC-I alleles. Here, we directly explored whether both house sparrow females and males could estimate by scent the number in MHC amino acid and functional variants as well as the level of MHC-I similarity or dissimilarity of potential partners. Our results show that neither females nor males exhibit a preference related to the number of MHC-I amino acid variants or functional variants or in relation to MHC amino acid or functional similarity of potential partners, suggesting that MHC-I is not detected through olfaction. Further studies are needed to understand the mechanisms responsible for MHC-I based mate discrimination in birds.

## Introduction

When selecting partners for mating, females may choose those that will provide the best resources, genes or both in order to increase offspring fitness [1, 2]. Most experimental studies addressing sexual selection in birds have focused on male quality to explain adaptive mate choice, considering females to assess potential mates by using male ornamental traits, such as plumage coloration [3, 4], or acoustic traits such as male songs [5], that are condition-dependent signals of male quality and therefore, may allow them to obtain resources and/or good genes for their offspring. In fact, in many passerine species, male parental care of offspring is needed to ensure nestling survival. In contrast, studies addressing the mate choice of females in relation to genetic dissimilarity are less well described. However, the increased use of molecular techniques for assessing paternity and genetic compatibility are providing examples for female mate choice that is not exclusively based on genes reflecting quality, but indeed suggest additional mechanism critical to female choosiness. Females may thus use alternative mechanisms for choosing mates such as extra-pair copulations [6, 7], or post-copulatory choice [8].

The study of alternative sexual selection is providing examples of major histocompatibility complex (MHC) dissimilar selection in birds ([7, 9, 10], but see [11]), pointing out that clutch sizes and the probability to extrapair relationships that can be affected by the partner’s MHC diversity [7, 9, 10]. For example, although blue tit *Cyanistes caeruleus* females are known to use song [12] and plumage coloration [13] when selecting mates, there is evidence that they select extrapair partners in based on their genetic dissimilarity, as extra-pair nestlings showed higher degree of heterozygosity compared to within-pair nestlings [14]. Since nestlings with higher degree of heterozygosity were more likely to survive [14], females may increase their fitness by selecting genetically dissimilar mates when engaging in extrapair copulations. Although, females may ensure all nestling survival by choosing the best parental male based on phenotypical characteristics such as plumage coloration [13], they may indeed use other characteristic male signals such as genetic heterozigosity [15, 16].

In order to provide the best genetic benefits to their offspring females can select males with “good genes”, that will consequently be inherited by their offspring and may increase their fitness [17] or select more heterozygous or dissimilar males to increase the genetic diversity of offspring [18]. When this heterozygosity is exhibited in genes of the immune system, the benefits for offspring fitness are obvious in terms of resistance to a wide range of infections [19]. The MHC is a large chromosomal region containing several highly polymorphic genes (MHC class I and II loci) that play a central role in controlling immunological self/non-self recognition [20, 21] and encode cell-surface glycoproteins that mediate antigen presentation to T-lymphocytes. Therefore, MHC genes play a fundamental role in immune recognition of pathogens and parasites. Specifically, MHC I genes are involved in recognition of intracellular pathogens whereas MHC II genes are involved in the recognition of extracellular parasites. For example, in house sparrows, *Passer domesticus*, specific MHC class I alleles are associated with stronger immune T cell responses to MHC I presented antigen [22] and increased resistance to malaria parasites [23, 24]. Furthermore, house sparrow females exhibiting lower numbers of functional MHC I alleles lay eggs with more eggshell bacteria [25], and some MHC genes have been found to partly explain early survival in this species [26]. Consequently, both pathogen-mediated selection [27] and sexual selection [28-31] may explain the maintenance of MHC polymorphism.

Although, the evidence is not as well described as in mammals [32, 33], including humans [34], evidence for MHC-associated mate choice has been found in fishes [35] and reptiles [36]. However, several observational studies in birds suggest that MHC genes play a role in mate choice ([7, 9, ], but see [11, 37]. While olfaction has been ascribed to account for MHC recognition in other taxa, sexual selection based on olfaction has formerly been neglected in avian studies. However, the study by Leclaire and colaborators [15] showed first evidence that birds can use olfactory cues to assess MHC similarity. Here incubating males preferred the scent of MHC II-dissimilar females, while incubating females preferred the scent of MHC-similar males [15]. Conversely, a study assessing the scent preferences in song sparrows *Melospiza melodia* showed that during breeding season both female and males spent longer time periods close to odours from MHC II-dissimilar partners [16]. Given these controversial results, further studies are needed to disentangle the MHC scent preferences of birds in the context of sexual selection during mating season. To our knowledge, MHC class I scent preferences of birds in the context of sexual selection has not been previously assessed.

In this study we have experimentally explored whether olfactory signals play a role in the preference for potential partners with greater MHC I dissimilarity and/or diversity in house sparrows, *Passer domesticus*.

## Materials and methods

### Study Species

The house sparrow was chosen as a model species as their olfactory capabilities have been previously described in the context of roosting cavity assessment [38] and social interactions [39]. Moreover, in the house sparrow, MHC class I allele diversity is associated with reproductive success [40]. It has been shown that males with low MHC diversity or too dissimilar MHC alleles fail to form breeding pairs [41]. Therefore, female house sparrows likely assess their male mate preferences based on their own MHC allelic diversity. Females with low MHC allelic diversity choose males with relatively higher MHC allelic diversity, whereas no preference is exhibited by females with high MHC diversity ([42], but see [43]).

We captured with mist nets 100 male and 51 female adult house sparrows *Passer domesticus* at the Madrid Zoo (Spain) during February and March 2015. Immediately after capturing, we measured birds with a dial calliper to the nearest 0.01 cm and birds were weighed with a spring balance to the nearest 0.1 g. We measured the length of the visible badge of males with a digital calliper to the nearest mm from the base of the bill to the point on the black breast at which the black feathers finished. This measurement was used instead of the area of the badge, because measurement of the width across the whole black area is less accurate, and it has been used as a measure of badge size in previous studies [44]. All birds were individually banded with numbered aluminium and PVC rings. We obtained blood (80-100 μl) from the brachial vein with the aid of a needle and a capillary tube. Blood samples were centrifuged (2000 × *g*, 5 min) with a portable centrifuge (Nahita 2507-01). Serum and cellular fractions were separated, and the cellular fraction was conserved in ethanol and maintained for later analysis.

Birds were housed at the FIEB, Foundation for the Research and the Study of Ethology and Biodiversity (Casarrubios del Monte, Toledo), in outdoor aviaries, until two weeks before the experiments, when birds were individually housed in cages inside the aviaries, so they were maintained at outdoor temperature and photoperiod. Bird weight and badge size was again measured at this time point.

### MHC Characterization

For DNA metabarcoding library preparation, PCRs were carried out in a final volume of 25 μl, containing 2.5 μl of template DNA, 0.5 μM of the primers (GCA21M, 5’ CGT ACA GCG GCT TGT TGG CTG TGA, and fA23M, 5’ GCG CTC CAG CTC CTT CTG CCC ATA, primers to which the Illumina sequencing primer sequences were attached to their 5’ ends), 12.5 μl of Phusion DNA polymerase mix (Thermo Fisher Scientific), and ultrapure water up to 25 μl. The reaction mixture was incubated as follows: an initial denaturation at 98 °C for 30 s, followed by 35 cycles of 98 °C for 10 s, 60 °C for 20 s, 72 °C for 20 s, and a final extension step at 72 °C for 10 minutes. The index sequences which are required for multiplexing different libraries in the same sequencing pool were attached in a second PCR round with identical conditions and only 5 cycles. The libraries were run on a 1 % agarose gel stained with REAL Safe (Durviz), and imaged under UV light. Negative controls that contained no DNA were included to check for contamination during library preparation. Libraries were pooled in equimolar amounts according to band intensity. The pool was purified using the Mag-Bind RXNPure Plus magnetics beads (Omega Biotek), following the instructions provided by the manufacturer. The pool was sequenced in a MiSeq PE300 run (Illumina).

The data were filtered to remove low-quality sequences, any reads representing artefactual MHC alleles or putatively non-functional sequences. Firstly, a 1% bioinformatics filter was applied. After that, a minimum total sequence abundance/individual was set to 200, accordingly to [37]. After data processing, we obtained 117 different MHC class I alleles in 151 individuals. The average (± SE) number of alleles per individual was 4.62 ± 0.15 (range: 1–10).

### Amino acid and functional distance between birds

Amino acid and functional distances between individual genotypes were used to describe MHC similarity between individuals and were calculated following the approaches described in [15, 16, 37]. To calculate amino acid distances, a maximum-likelihood tree was inferred for all translated MHC sequences, using PhyML (v. 3.0) [45] in the ATGC platform with a JTT substitution model, selected automatically by SMS [46], following the Bayesian Information Criterion. Because this metric can be sensitive to cutoff thresholds and other methodological decisions [47, 48], in addition to calculating distances for the full data set, we also generated four additional phylogenies, each removing one of the four alleles with the longest branch lengths. We calculated unweighted UniFrac distances for each of the 5 phylogenies, then used the average of all analyses [16].

To calculate functional distances, we obtained the functional MHC alleles from Lukasch et al. [26], that are based on the chemical binding properties of the amino acids in the peptide binding regions (PBRs), described by five physio-chemical descriptor variables (z-descriptors) for each amino acid [49]. We obtained 113 functional MHC class I alleles. The average (± SE) number of functional alleles per individual was 4.48 ± 0.15 (range: 1–10). The functional MHC alleles obtained are shown in Table 2 (supplementary material). The resulting matrix of allele frequencies was used to construct an alternative maximum-likelihood tree with contml in the PHYLIP-package, v. 3.695. This tree represents clusters of functionally-similar MHC sequences rather than clusters of evolutionary-similar MHC sequences. The amino acid and functional trees were used as references from which the amino acid and functional distances between MHC-sequence repertoires were calculated, using unweighted UniFrac analyses (Phyloseq v. 1.22.3 package in R). We also generated seven additional phylogenies, each removing one of the seven functional alleles with the longest branch lengths, and calculated mean unweighted UniFrac distances across all eight functional trees. Amino acid distances were positively correlated with functional distances among the 151 individuals used in this study (Mantel test: *r* = 0.87, *P* = 0.01; calculated using Vegan v. 2.4-6 package in R).

### Behavioural Experiment

The experiment was performed indoors in May and June 2015 using an olfactometry chamber (Fig 1). The experimental device was composed of a small central plastic box (15 × 25 × 25 cm) where the experimental bird was placed. The chamber contained a small 12 V PC fan that extracted the air from the device creating a low-noise-controlled airflow (Fig 1). In each test, a bird was introduced into the central box and maintained in the dark for 5 min after that the doors were opened. Each choice chamber was divided into two sectors with screens. The distal sectors of the choice chambers (15 × 25 × 25 cm) contained two little cages where the scent donor birds were situated. Both, the doors connecting the central chamber with the choice chambers and the screens creating the sectors, were made with a dense plastic mesh that allowed air flow but prevented birds from seeing through them. The device was sealed and only openings at the farthest walls of the choice chambers allowed air flow. The fan created two constant air flows, each one entering across the openings located at the farthest walls of each choice chamber, passing by the donor birds and crossing the central chamber, and going outside from the device through the fan. Thus, the focal bird received two separate air flows, carrying the scent of the corresponding donor bird. Donor birds were temporarily kept in darkness and reduced space, preventing them from moving or calling. Therefore, the experimental bird only perceived the scent of the donor birds without visual or acoustics contact. The experimental room was to the greatest possible extend sealed from exterior noise, enabling the experimenter to perceive any acoustic signals from any of the birds in the device. The device and the methodology have been successfully used in social contexts before [50, 51].

**Fig 1.**
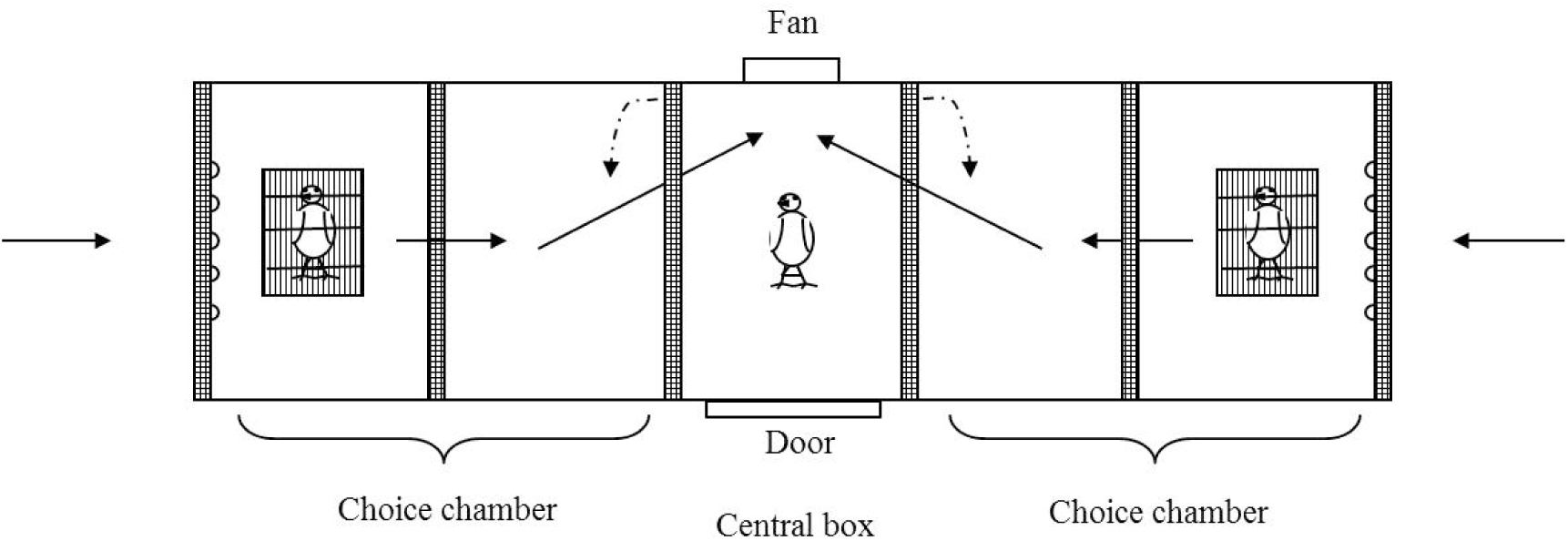
**Olfactometry chamber. The solid arrows indicate the direction of air flow within the chamber, whereas the dashed lines indicate the direction of opening of the two doors connected to the two choice plastic chambers**.

We noted down the choice chamber that was first approached by each tested focal bird after opening the central chamber. The assessment of “first choice” as a preference measure of birds to chemical stimuli has been previously demonstrated [15, 50, 51]. To minimize the experimental time duration of the trials and to release the birds as soon as possible, birds that did not show any choice preference after 1 min, were motivated to make an active choice by gently knocking on the middle of the entry door of the central chamber to motivate the bird to move to any of the choice chambers. The knocking on the door did not influence the preference of birds (see results).

We selected donor birds of the same sex, similar body size and body condition. When using male donor birds, we chose birds with similar badge size. Therefore, we offered focal birds the scent of two potential mates of similar body condition and dominance status in the case of males. There were no significant differences in the body condition (repeated measured ANOVA, *F*_1,127_ = 0.60, *P* = 0.44), or the badge size (*F*_1,48_ = 0.09, *P* = 0.77) between both scent donor birds. The MHC characterization was done a posteriori. We tested 72 males and 48 females. Only one female bird did not move to any chamber even after knocking on the door, and was therefore excluded from the analysis. We used 68 different pairs of scent-donor birds, each pair was used on average 1.76 (± 0.8) times. Focal birds made 61 % of their first choices to the left and 39 % of the choices to the right side. Birds were returned to their cages as soon as they were tested. The olfactometry device was carefully cleaned with alcohol in between trials and experiments continued after the alcohol was completely evaporated.

### Data Analysis

To analyse whether birds could detect amino acid variant numbers of potential mates by using chemical cues alone, we performed a generalized linear mixed model with binomial errors and a logit link function (GLMM). We modelled the probability that birds chose the scent of the conspecific of the opposite sex with greater number of MHC-I amino acid variants (as a dichotomous variable: greater number of MHC amino acid variants (yes) vs. lower number of MHC amino acid variants (not)) in relation to the sex and the number of MHC amino acid variants of the focal bird. We included the interaction between the sex and number of MHC amino acids of the focal bird in the model to test whether males and females differed in their preferences for partners with different numbers of MHC amino acid variants in relation to their own number of MHC amino acid variants. We also included the pair of donor birds in the model as a random factor to control for the fact that pairs of donors could be used more than once. We included in the initial model a variable reflecting whether the experimental bird left the chamber when we opened the doors or after 1 min as fixed factor.

We performed a similar analysis to assess whether birds chose the scent of the conspecific of the opposite sex with greater number of MHC-I functional alleles, including in number of MHC-I functional alleles of the focal bird in the model, the sex and the interaction between the number of MHC-I functional alleles and the sex in the model. We also included the pair of donor birds in the model as a random factor. We included in the initial model a variable reflecting whether the experimental bird left the chamber when we opened the doors or after 1 min as fixed factor.

We also performed two generalized linear mixed models with binomial errors and a logit link function (GLMM) to model the probability that birds chose the scent of the conspecific of the opposite sex with greater a) MHC-I amino acid distance and b) MHC-I functional distance (as a dichotomous variable: greater MHC-I amino acid or functional distance (yes) vs. lower amino acid or functional distance (not)) in relation to the sex of the focal bird. We included in the initial model a variable reflecting whether the experimental bird left the chamber when we opened the doors or after 1 min as fixed factor. We also included the pair of donor birds in the model as a random factor. Data analyses were performed with R program [52].

### Ethical Note

After capture, birds were housed in ten aviaries (2.5 ×2.5 × 2.5 m), separated by sex (10 females per aviary and 16 males per aviary). Aviaries contained vegetation (bamboo branches) that birds could use as perches, and grass and sand on the ground. Commercial food for granivorous passerines and water were provided *ad libitum*. In order to reduce the time of capturing individual birds used within the trials, two weeks before the experiments, birds were individually housed in cages (60 × 40 × 40 cm) inside the aviaries, so they were maintained at outdoor temperature and photoperiod. After the behavioural tests were completed, birds were released again in the aviaries for two weeks and thereafter released at their capture site. Birds maintained healthy throughout the experiments. Experiments were carried out under license of the Ethical Committee of the CSIC (268/2015), the Animal Experimental Committee of the Junta de Castilla la Mancha (202121) and the department of Flora and Fauna of Comunidad de Madrid (10/238509.9/14).

## Results

Our results do not show any preference of birds for the scent of potential partners having a greater number of MHC-I amino acid variants (*Z* = 1.70, *P* = 0.09, Table 1, Fig 2a) or MHC functional alleles (*Z* = 1.60, *P* = 0.11, Table 2, Fig 2b). There was not effect of sex in this lack of preferences (*Z* = -0.99, *P* = 0.32, and *Z* = -0.21, *P* = 0.83, respectively), and the interaction between the sex and the number of amino acid variants or functional alleles was not significant in either analysis (*Z* = 0.83, *P* = 0.41 and *Z* = 0.37, *P* = 0.71, respectively). Neither the number of MHC amino acid variants (*Z* = -1.59, *P* = 0.11) nor the number of functional alleles (*Z* = -1.47, *P* = 0.14) influenced bird choice. Whether the experimental bird left the chamber when opened or after 1 min as fixed factor did not affect the choice of the respective bird (see Table 1 and 2).

**Table 1.** **Model for bird choice in relation to the number of MHC-I amino acid variants of a potential partner (as a dichotomous variable: greater number of MHC amino acid variants (yes) vs. lower number of MHC amino acid variants), in relation to the sex and the number of MHC amino acid variants of the focal bird, the interaction between the sex and number of MHC amino acids of the focal bird, the variable reflecting whether the experimental bird left the chamber when we opened the doors or after 1 min as fixed factor and the pair of donor birds as a random factor**.

|  | Estimate | SE | Z | P |
| --- | --- | --- | --- | --- |
| Intercept | 1.44 | 0.85 | 1.70 | 0.09 |
| Sex | -1.30 | 1.32 | -0.99 | 0.32 |
| Choice within 1 min or after | -0.06 | 0.54 | -0.11 | 0.91 |
| Number of MHC amino acid variants | -0.28 | 0.17 | -1.59 | 0.11 |
| Sex x Number of MHC amino acid variants | 0.21 | 0.26 | 0.83 | 0.41 |

**Table 2.** **Model for bird choice in relation to the number of MHC-I functional alleles of a potential partner (as a dichotomous variable: greater number of functional alleles (yes) vs. lower number of functional alleles), in relation to the sex and the number of MHC functional alleles of the focal bird, the interaction between the sex and number of functional alleles of the focal bird, the variable reflecting whether the experimental bird left the chamber when we opened the doors or after 1 min as fixed factor and the pair of donor birds as a random factor**.

|  | Estimate | SE | Z | P |
| --- | --- | --- | --- | --- |
| Intercept | 1.44 | 0.90 | 1.60 | 0.11 |
| Sex | -0.31 | 1.44 | -0.21 | 0.83 |
| Choice within 1 min or after | -0.22 | 0.54 | -0.41 | 0.68 |
| Number of MHC functional alleles | -0.28 | 0.19 | -1.47 | 0.14 |
| Sex x Number of MHC functional alleles | 0.10 | 0.28 | 0.37 | 0.71 |

**Fig 2.**
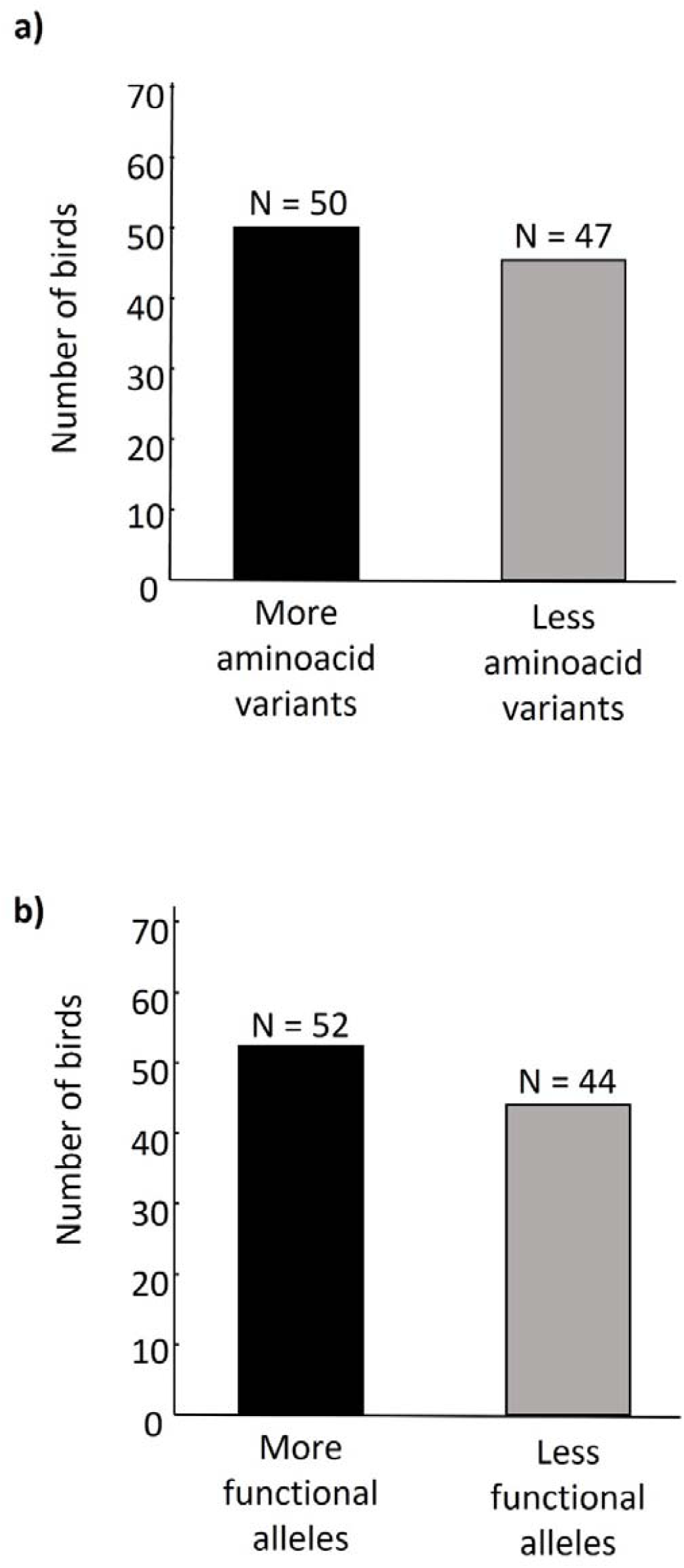
Number of birds that chose the side of the olfactometry chamber containing the scent of a conspecific of the opposite sex with greater (black) or lower (grey) a) number of MHC-I amino acid variants or b) number of MHC-I functional variants.

Our results do not show any preference of birds for the scent of potential partners having a greater MHC-I amino acid distance (*Z* = 0.27, *P* = 0.64, Table 3, Fig 3a) or functional distance (*Z* = 0.28, *P* = 0.35, Table 4, Fig 3b). Neither the sex (*Z* = 0.38, *P* = 0.70, and *Z* = 0.96, *P* = 0.33, respectively) nor whether the bird made the choice within the first minute or afterwards (*Z* = 0.42, *P* = 0.74, and *Z* = -1.08, *P* = 0.28, respectively) influenced their choice.

**Table 3.** **Model for bird choice in relation to the difference in the MHC-I amino acid distance between the focal birds and the two scent donor birds of the opposite sex (as a dichotomous variable: greater MHC-I amino acid distance (yes) vs. lower MHC-I amino acid distance (no)) in relation to the sex of the focal bird, the variable reflecting whether the experimental bird left the chamber when we opened the doors or after 1 min as fixed factor and the pair of donor birds as a random factor**.

|  | Estimate | SE | Z | P |
| --- | --- | --- | --- | --- |
| Intercept | -0.12 | 0.27 | -0.46 | 0.65 |
| Sex | -0.15 | 0.38 | -0.39 | 0.70 |
| Choice within 1 minute or after | -0.14 | 0.42 | -0.33 | 0.74 |

**Table 4.** **Model for bird choice in relation to the difference in the MHC-I functional distance between the focal birds and the two scent donor birds of the opposite sex (as a dichotomous variable: greater MHC-I functional distance (yes) vs. lower MHC-I functional distance (no)) in relation to the sex of the focal bird, the variable reflecting whether the experimental bird left the chamber when we opened the doors or after 1 min as fixed factor and the pair of donor birds as a random factor**.

|  | Estimate | SE | Z | P |
| --- | --- | --- | --- | --- |
| Intercept | 0.26 | 0.27 | 0.94 | 0.35 |
| Sex | -0.38 | 0.38 | -0.99 | 0.33 |
| Choice within 1 minute or after | -0.45 | 0.42 | -1.08 | 0.28 |

**Fig 3.**
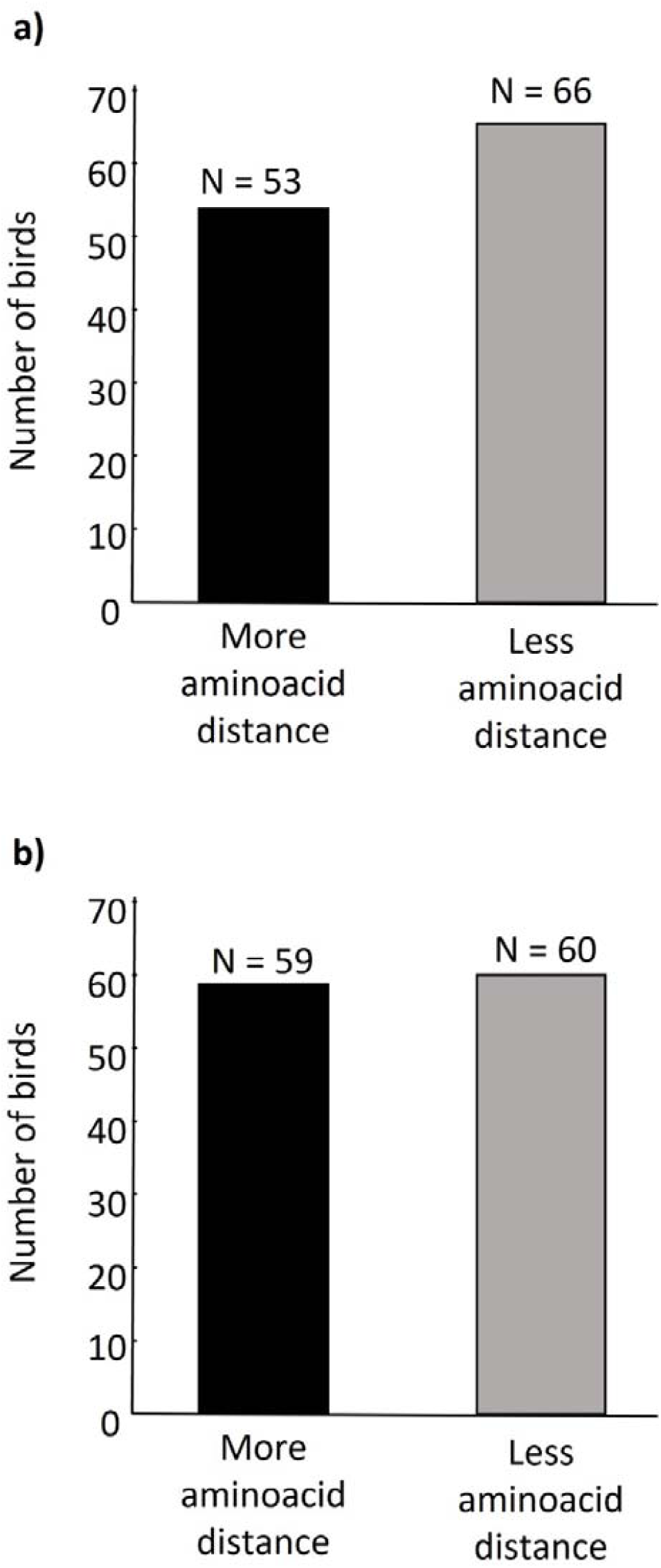
Number of birds that chose the side of the olfactometry chamber containing the scent of a conspecific of the opposite sex with greater (black) or lower (grey) a) MHC-I amino acid distance or b) MHC-I functional distance.

## Discussion

Our experimentally obtained results on house sparrows do not provide evidence of MHC-I scent discrimination in a passerine bird species. Although olfaction has been demonstrated to play a major role in assessing genetic dissimilarity in birds [15, 16] as well as in other taxa [53], our results, however, show that neither females, nor males, exhibited any preference for the scent of conspecifics with greater allelic diversity or more MHC-dissimilarity. This is contrary to previous studies showing that female blue petrels *Halobaena caerulea*, preferred the scent of MHC-similar males [15], and female song sparrow *Melospiza melodia* preferred the scent of MHC-dissimilar and more MHC-diverse males during the mating period [16]. Differences between our results and those of Grieves and collaborators could be due to the different methodology used to test bird scent preferences as well as to the level of sociality of both study species. Grieves and collaborators analysed the percentage of time spent close to the two stimuli after 5 minutes from the beginning of the trial. Neither the first choice nor the preference of birds during the first 5 minutes of the trial was reported. We in contrast, analysed the first choice of birds. The validity of first choice as a measure of the interest of birds in particular chemical stimuli has been previously demonstrated [50, 51, 54, 55, 56], including MHC related scents [15]. Furthermore, our results are robust because we used a large sample size used and because we used scent-donor birds of similar body condition in each trial to exclude any potential effect of body condition of scent-donor birds in the choice of focal birds.

While there is no reason to expect that all bird species use the same type of information to evaluate potential partners, nevertheless our results are difficult to explain in a sexual context, because in house sparrows MHC class I diversity is associated with reproductive success [40]. It is also known that males with low degree of MHC class I diversity or too dissimilar MHC alleles fail to form breeding pairs [41]. Furthermore, results of a previous study found that house sparrow females with low MHC allelic diversity choose males with high MHC allelic diversity [42], suggesting that house sparrow females show mate preferences based on the MHC of potential partners ([42], but see [43]). First choice is a good measure of the spontaneous interest of an animal to a particular cue, whereas time spent close to the stimulus [57] may rather be related to the behaviour shown later on in the series of events to a certain cue. Using life birds as scent donors, the first choice was a valid measure of the response of birds to scent in our study. To analyse whether the lack of preferences for the scent of most MHC class I dissimilar conspecifics is maintained over the time, more studies are needed to assess the subsequent response of birds to the scent of potential partners, for instance using uropygial gland secretions as scent sources.

Although several hypotheses have been proposed to explain the underlying mechanisms [53] is still not completely understood how MHC genes influence scent [58, 59]. As MHC proteins appear in urine and sweat, it was proposed that these molecules may constitute odorants [60]. However, since MHC proteins are non-volatiles this hypothesis is rather weak [61]. Another hypothesis proposes that the peptides bound by MHC molecules are the precursors of volatile odorants [62], or that those peptides can be detected by the vomeronasal organ [63]. Furthermore, it has been proposed that MHC genes may influence the bacterial profile that influences the scent of an individual [64, 65, 66, 67]. In many avian species, the source of scent that birds are detecting usually comes from externally applied secretions of the uropygial gland [68]. The quantity and composition of this secretion varies between species, sexes, ages, seasons, and is dependent on diet and hormonal levels [69]. This secretion conveys information on genetic compatibility [15,16], which may play a role during kin recognition [70] and mate choice [15, 16, 37].

Evidence that birds are using the volatile compound of uropygial gland secretions for detecting the MHC characteristics of conspecifics [16] comes from studies in song sparrows that were able to assess MHC similarity when exposed only to the uropygial gland secretion. However, MHC-related scent could also be produced by symbiotic bacteria [71] whose diversity is associated with MHC-II diversity [66]. Further research is needed to elucidate the source of MHC-related scent in birds.

In conclusion, the observed lack of preference for the scent of MHC dissimilar potential partners suggests that the assessment of genetic diversity and dissimilarity in MHC-I is not based on olfaction in our study population of house sparrows. However, our results do not fully exclude the possibility that mate choice in house sparrows includes the selection of partners based on the degree of similarity or dissimilarity in MHC genes. Here, increased MHC allele diversity in the progeny, resulting in higher resistance to pathogens [22] such as malaria parasites [23, 24, 72] may add to increased reproductive success [40]. We suggest that different methodological approaches in future investigation are needed to further the understanding of the mechanisms responsible for MHC-I based mate discrimination in birds.

## Acknowledgements

We especially thank Pilar Ochoa for laboratory analysis, Agustín López Goya and the personnel of Madrid Zoo for allowing the capture of birds in the Zoo. LA was supported by the Ramón y Cajal program. The study was funded by the Volkswagen Foundation (85 994-1). Authors declare no conflict of interest.

## Data Accessibility

All data shown/used are presented in this manuscript.

